# Mycorrhiza Co-Association with *Aspilia pruliseta* Schweif and Phosphorus Uptake Effects on Growth Attributes of Gadam Sorghum in Selected Sites in Kenya

**DOI:** 10.1101/2019.12.19.882100

**Authors:** J.P. Muchoka, D.N Mugendi, P.N Njiruh, C.N. Onyari, P.K. Mbugua, E.M. Njeru

## Abstract

Mycorrhiza fungi are important components of soil microbiota in the rhizosphere and greatly influence uptake of mineral elements to plants. A green house experiment was conducted at the University of Embu. The experiment involved use of sterilized polythene potting material sized 30 cm by 40 cm. The pots were filled two thirds the height of the potting material with soil from a predetermined source in Gakurungu, Tunyai and Kanyuombora in the upper eastern region in Kenya. The soil used in the pots was collected from the rhizosphere of *Aspilia pruliseta* Schweif vegetation as well as adjacent areas without this vegetation as a control at 0-20 cm, 21-40 cm and 41-60 cm for each of the soil types (silty clay, silt loam and sandy loam) used in the experiment. Two sorghum seeds inoculated with mycorrhiza fungi were planted in each pot and a similar number of pots planted with un inoculated sorghum seeds as a control. Each of the 4 treatments mentioned above, was replicated four times giving n=144. Each pot was watered after every two days using a two-litre watering can for the first one week. Thereafter, watering regime was reduced to once a week but ensuring the pots remained moist. Watering was done uniformly to all the pots. This was maintained for a period of thirty five days. Data was analysed using SAS edition 8.2. Seed emergence, hypocotyl development and stand count were enhanced at P≤0.05 in both mycorrhiza fungi inoculated gadam sorghum seeds and in pots whose soils were taken from the rhizosphere of *Aspilia pruliseta* plants. The growth attributes had a positive correlation to yield at 95% confidence. Soil phosphate level was enhanced in both cases of gadam seed inoculation with mycorrhiza and in soils previously grown *Aspilia pruliseta* vegetation.

## 1. Introduction

Use of Fertilizers to supplement soil nutrients in promoting plant growth is prevalent in modern agriculture [1]. Phosphorus (P) is the most important nutrient element (after nitrogen) limiting agricultural production in most regions of the world [2]. In East Africa, Phosphorus deficiency occurs in many soils, not only due to P depletion through crop harvest and erosion but also due to P-fixing soils in the region [3]. It is hypothesized that a complex association occurs between mycorrhiza, soils and plants, leading to availability of phosphorus in the soils in usable forms to plants. Virtually all crop plants (except Brassicaceae, Amaranthaceae and Polygonaceae) worldwide are host to some form of mycorrhizal association [4–5]

Among the three major nutrients (nitrogen, phosphorus and potassium) required by plants, phosphorus constitutes a particularly critical component because on one hand it is limiting for crop yield on a large proportion of global arable land and, on the other hand, it is a non-renewable resource [6–8]. Agricultural experts points to a phosphate crisis in the foreseeable future [9]. Different approaches have to be explored to ensure continued supply of phosphates in the soil and hence sustained food production. Soil biology has emerged over the last decade as a critical part of the knowledge base for successful and sustainable agricultural production [10]. A key component of this subject is the plant/mycorrhizal fungi relationship, which has enormous potential for improved management of contemporary farming systems [10–11].

Majority of terrestrial plant species are capable of interacting with mycorrhiza fungi (MF) in nature to provide an effective pathway by which phosphorus is scavenged and rapidly delivered to cortical cells within the root [12–13]. MF acts as a catalyst to concentrate P in a manner that makes it available to plants [14–16]. The plant supplies the fungi with sugars produced by photosynthesis, while the hyphae network improves the plant capacity to absorb water and nutrients [6, 17].

Mycorrhizal symbioses contribute significantly to plant nutrition and particularly to phosphorus uptake [18]. The large surface area of the mycorrhiza fungi hyphal network is very efficient in nutrient uptake [6, 19]. Furthermore, phosphorus is a highly immobile element because it is easily absorbed by soil particles and a phosphate free zone rapidly occurs around plant roots [20]. Extra radical hyphae extend beyond this depletion zone, absorbing bio-available phosphate that would otherwise not be accessible to the plant [21–22]. Phosphate ions in soil also become rapidly bound with cations, forming insoluble complexes that are unavailable to plants [21–22].

Recent physiological and molecular research has revealed that the MF pathway plays a major role in P uptake, regardless of the extent to which a mycorrhiza fungi plant benefits in terms of increased growth or P uptake [23]. This experiment investigated *Aspilia pruliseta* Schweif plant in mediating phosphorus mining from the soil aided by mycorrhiza and eventual phosphates availability in soil colloids for use by other plants. Performance trials were conducted with Gadam sorghum (*Sorghum bicolor* L.) grown on rhizosphere soils of *Aspilia pruliseta* and growth attributes observed over time.

## 2.0 Materials and Methods

### 2.1 Study area

This experiment was set up at the University of Embu located on latitude 00 31’ 52.03” N and longitude 37 27’ 2.20” E at an elevation of 1480 m above sea level. The average annual rainfall around the university green houses is 1252 mm per year and is received in two distinct rainy seasons; the long rains (mid-March to June) with an average rainfall of 650 mm and the short rains (mid- October to December) with an average of 450 mm (Jaetzold et. al. 2006). The area has a mean annual temperature of 19.5°C, a mean maximum of 25°C, and a mean minimum of 14°C. The soils are mainly humic nitisols. Over 65% of the rains occur in the long rain season [24]. The soils are mainly humic nitisols derived from basic volcanic rocks [24], which are deep, well weathered with moderate to high inherent fertility but over time soil fertility has declined due to continuous mining of nutrients without adequate replenishment.

### 2.2 Methods

Mature plots of *Aspilia pruliseta* vegetation were identified in preselected sites in Tharaka Nithi (Gakurungu at 00°12’00”S, 37°51’00”E and Tunyai at 00°10’00”S, 37°50’00”E) and Embu (Kanyuombora at 0°21’0”S, 37°28’30”E) counties. The specific sites where soil was extracted were purposively selected and each sub site had a colony of *Aspilia pruliseta* vegetation, silty clay, silt loam and sandy loam soils. Soil categorization was based on USDA soil taxonomy. A five metre length tape measure was used to delineate the clearance area of 1m^2^ of the selected sub site. The underneath vegetation was cleared using a panga. Using a mattock, a forked jembe and a spade about 20 kgs soil from each depth of interest was dug in each of the sub sites at the depths of 0-20 cm, 21-40 cm and 41-60 cm. A three metre tape measure was used to take the vertical distances. The soil was put in separate, sterilized hessian bags that were clearly labeled using a felt pen for each depth of soil. Similar procedure was used to collect soil in the sites and sub sites for the control experiment but in adjacent areas where *Aspilia* was not growing. All other parameters remained the same, except the lack of *Aspilia pruliseta* vegetation in the patches selected. The soil collected was transported in clearly labeled hessian bags to the University of Embu green house using a three tone truck. At the green house, the soil was thoroughly mixed for each respective soil type, depth and presence or absence of *Aspilia* vegetation. Further to mixing the soil as above, soil was filled one third full into 30 cm by 40 cm sterilized pots. Two seeds of mycorrhiza fungi-inoculated Gadam sorghum [25] were planted into pots and a similar number planted with un inoculated sorghum seeds as a control. Each of the 4 treatments (Figure 1) as mentioned was replicated four times, giving n=144. Each pot was watered after every two days using a two-litre watering can for the first one week. Thereafter, watering regime was reduced to once a week but ensuring the pots remained moist. Watering was done uniformly to all the pots. This was maintained for a period of thirty five days. Data on plant growth attributes was taken every week and corroborated with treatments given in the pots.

**Figure 1:**
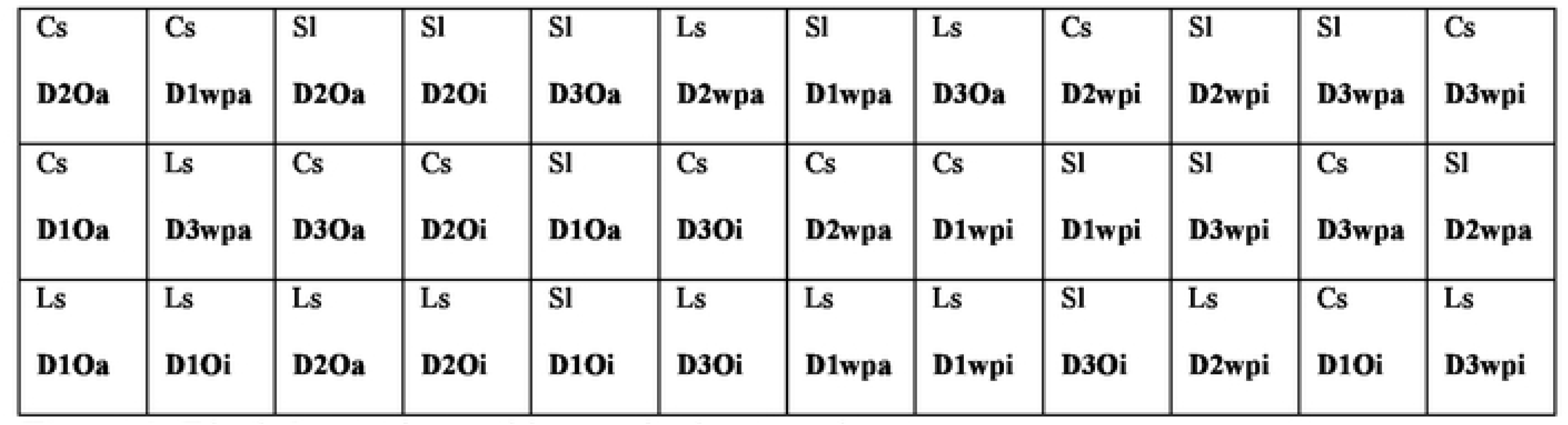
Block I experimental lay out in the green house **Legend:** SI-sandy loam soil; Cs-silty clay soil; Ls-silt loam soil; 0-soils without *Aspilia pruliseta* vegetation; wp soils with *Aspilia prulisela* vegetation; a Gadam seeds not inoculated and i Gadam seeds inoculated.

### 2.3 Data Analysis Methods

Data obtained from various treatments was subjected to SAS Edition 8.2. Differences between treatments means was examined using least significant difference (LSD) at P ≤0.05. Plant growth attributes obtained on the soil types, rhizosphere depths and locations with *Aspilia pruliseta* were compared to their controls and conclusions made.

## 3. Results

### 3.1 Effect of soils on gadam sorghum seed emergence and stand county

Soil type influenced gadam sorghum seeds emergence. Greater seed emergence was experienced in silt loams at 97% within the first seven days. Seeds planted in silty clay soils had the lowest seed emergence in the first week and significantly differed from the other soils at P≤0.05 (Table 1)

**Table 1:** Percentage stand count of gadam sorghum in 3 main soil types for the first 35 days of growth

| Factor | SC7 | SC14 | SC21 | SC28 | SC35 |
| --- | --- | --- | --- | --- | --- |
| Silty Clay | 83.3b | 83.3b | 83.3b | 84.4b | 82.3b |
| Silt Loam | 96.9a | 96.9a | 94.8a | 94.8a | 94.8a |
| Sandy Loam | 90.2ab | 90.2ab | 90.2ab | 90.2ab | 90.2a |
| LSD | 7.2 | 7.2 | 7.6 | 7.6 | 7.8 |

SC-Stand count, after 7 days, 14 days, 21 days and 35 days

### 3.2 *Aspilia pruliseta* effect on gadam sorghum seeds early growth

Gadam seed emergence and early growth was highest in soils that had *Aspilia pruliseta* vegetation (Figure 2) at 96.5% in the first week and differed significantly with seeds planted in soils that were not previously growing *Aspilia pruliseta* at P≤0.05 (Table 2). Similarly, stand count was almost maintained in soils with *Aspilia pruliseta* vegetation than those without (Figure 2).

**Table 2:** Percentage stand count of gadam sorghum seedlings on *Aspilia* soils and those without

| Factor | SC7 | SC14 | SC21 | SC28 | SC35 | SC42 | SC49 | SC56 | SC63 | SC70 | SC77 | SC84 |
| --- | --- | --- | --- | --- | --- | --- | --- | --- | --- | --- | --- | --- |
| Without Ap | 83.8b | 83.8b | 83.8b | 84.4b | 83.1b | 79.4b | 78.6b | 78.3b | 77.6b | 77.3b | 77.3b | 77.3b |
| With Ap | 96.5a | 96.5a | 95.1a | 95.1a | 95.1a | 94.4a | 95.1a | 95.1a | 95.1a | 95.1a | 95.1a | 95.1a |
| LSD | 5.9 | 5.9 | 6.2 | 6.2 | 6.3 | 7.2 | 7.2 | 7.3 | 7.3 | 7.4 | 7.4 | 7.4 |

SC-stand count 7days to stand count at day 84 (harvesting)

**Figure 2:**
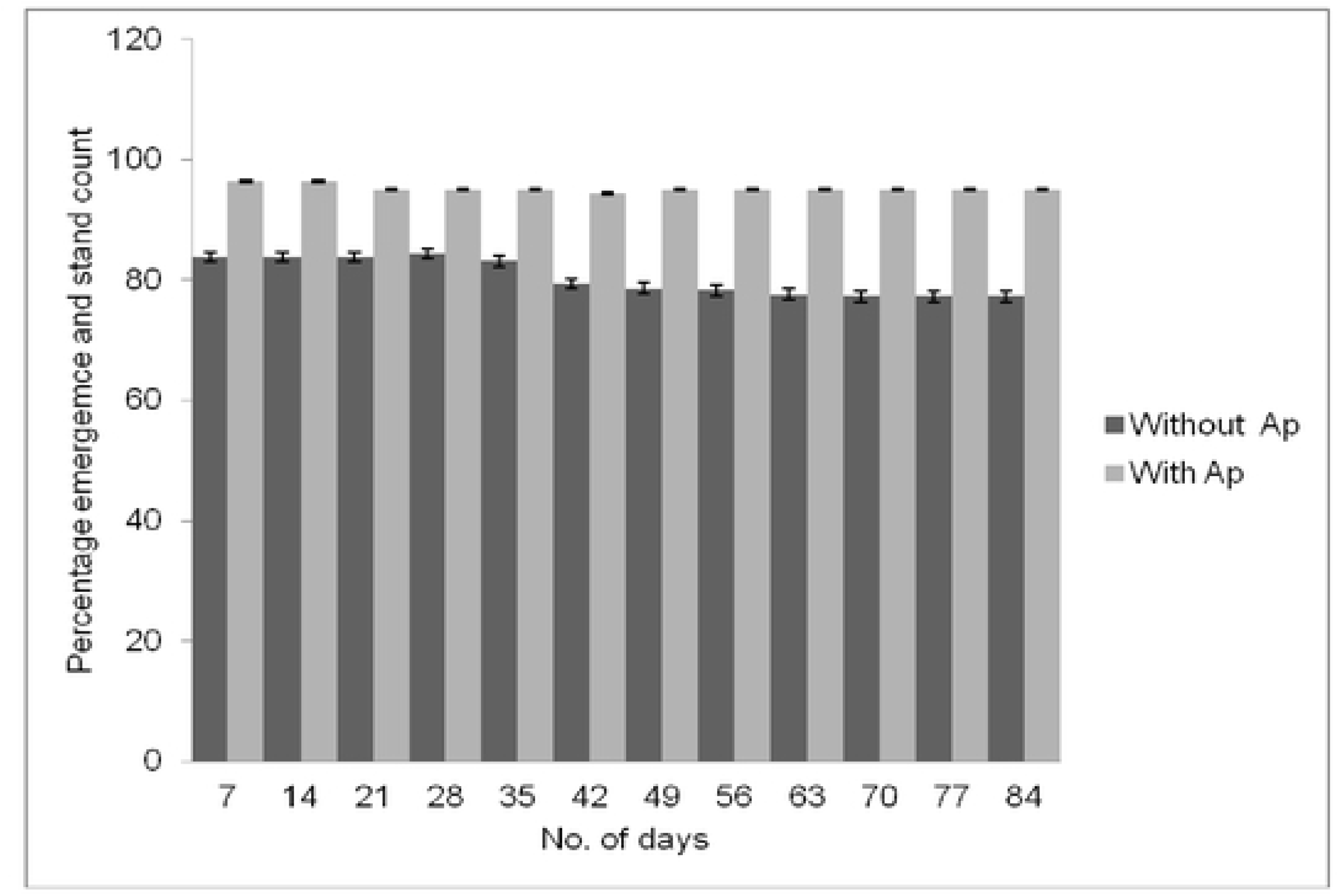
Percentage gadam seed emergence and stand count over the growing period

### 3.3 Effect of gadam sorghum seeds inoculation with mycorrhiza fungi on seed emergence, stand count and plant height

Gadam seed emergence and stand count gave similar results when the sorghum seeds were inoculated with mycorrhiza fungi (Figure 3). The inoculated seeds had higher per cent seed emergence and stand count compared to the un inoculated. In addition, aggregate gadam crop stability was higher in inoculated sorghum seeds (Figure 3).

**Figure 3.**
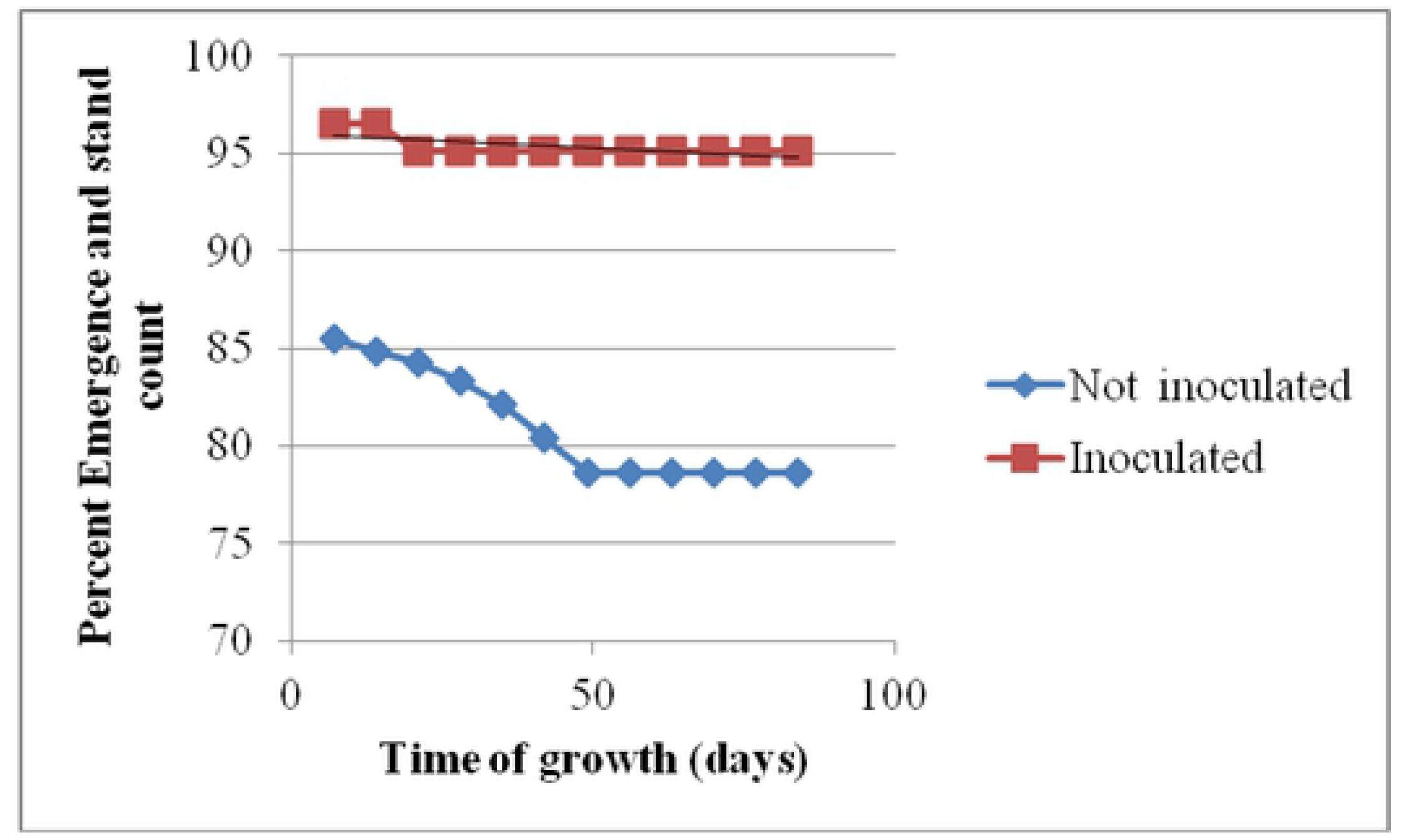
Sorghun1 emergence and stand count for inoculated and un inoculated seedlings

The average sorghum plant height was 124 cm at physiological maturity (Figure 4). Although both growth curves were normal, the curve with soils where *Aspilia pruliseta* previously grew peaked higher. Similar results were obtained when gadam sorghum seeds were inoculated with mycorrhiza fungi from the rhizosphere of *Aspilia pruliseta.*

**Figure 4:**
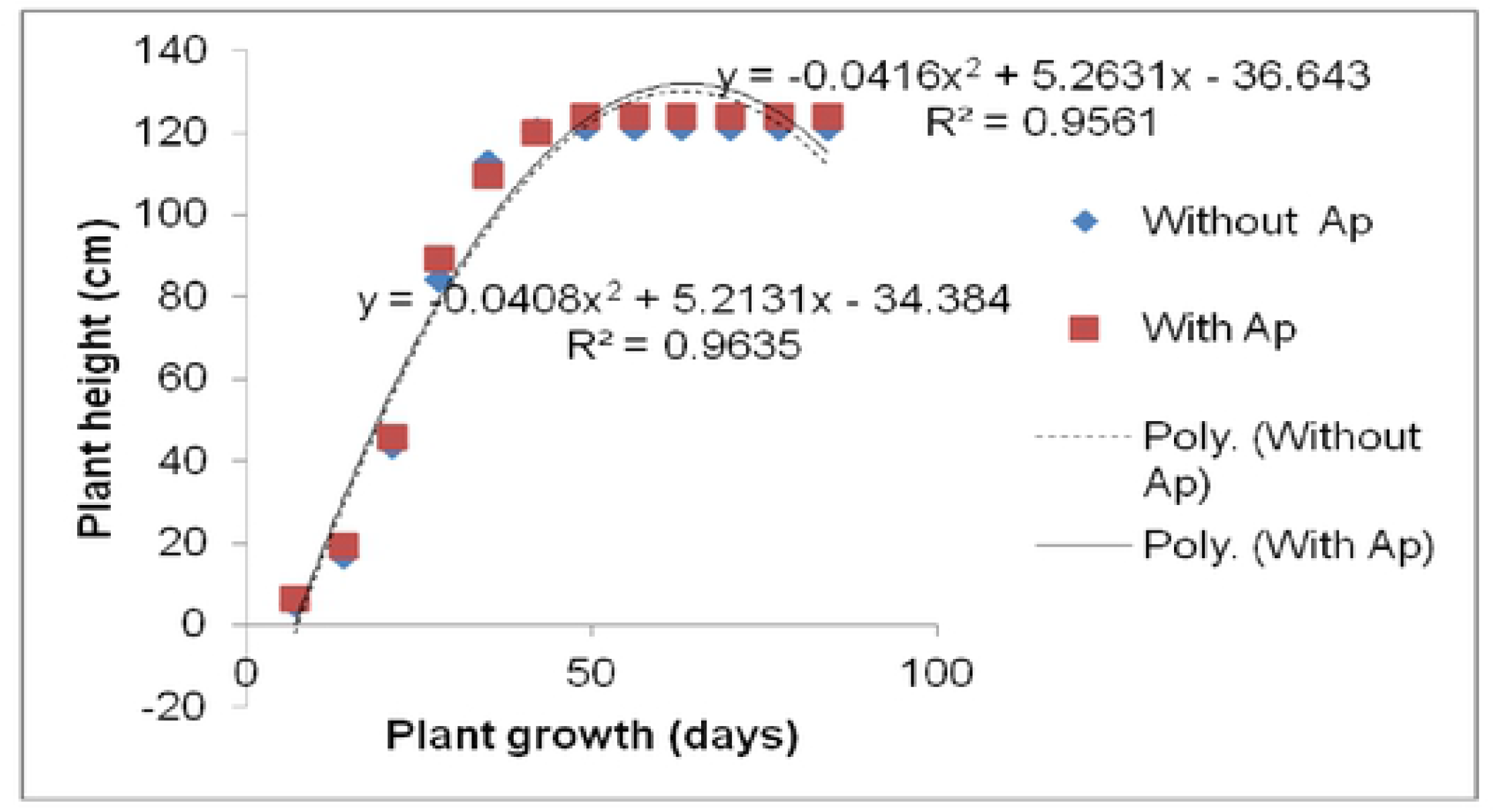
Gadam sorghum height over period *Aspilia* soils and non Aspilia soils

### 3.4 Changes in pH, phosphorus and phosphate levels before and after gadam sorghum harvest

The initial rhizosphere reading for pH and phosphorus was higher but the opposite was true of phosphates after the experiment (Table 3). Phosphate level in soils grown with *Aspilia pruliseta* vegetation had a higher content compared to soils without the vegetation (Table 3)

**Table 3:**
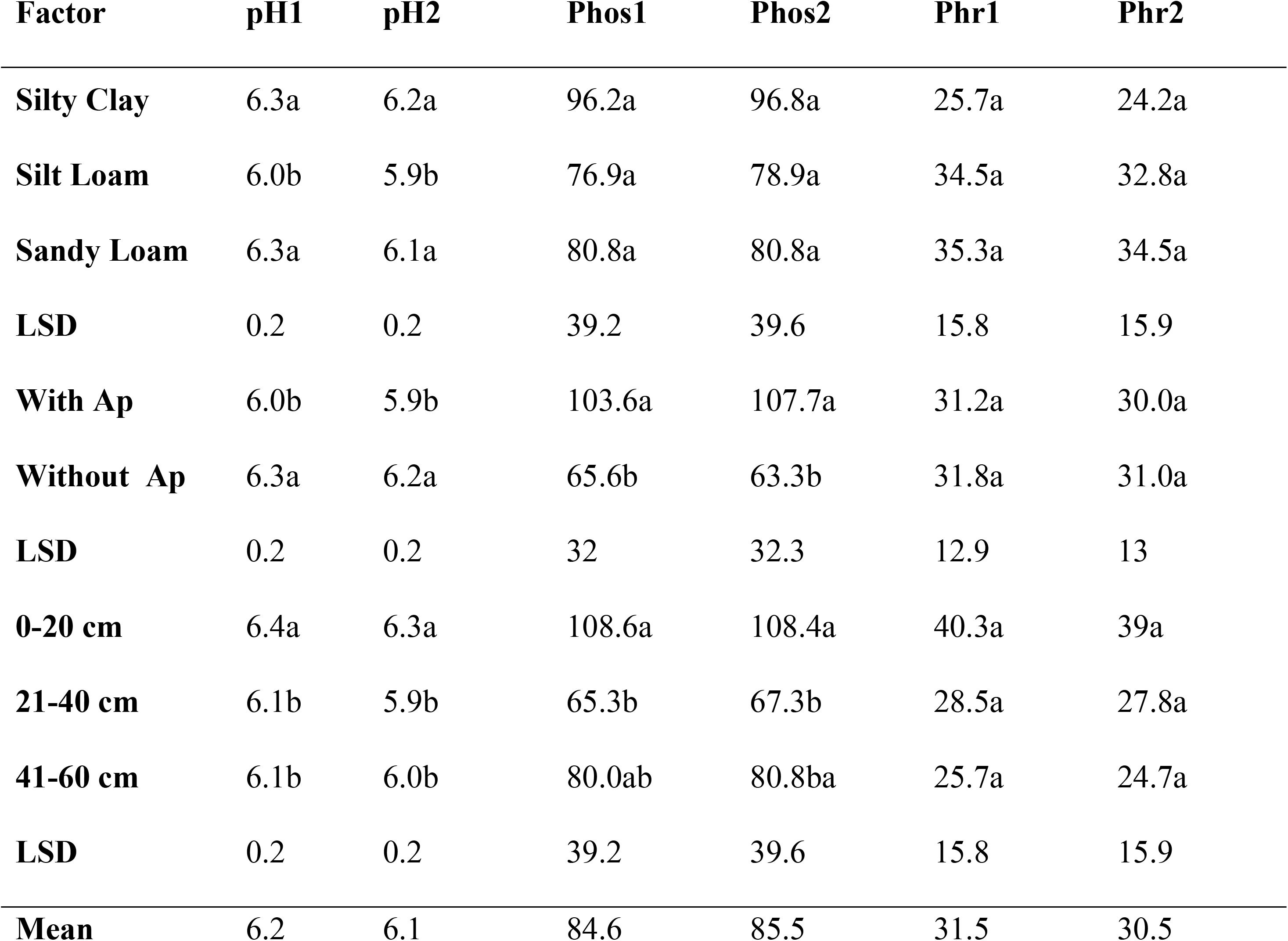
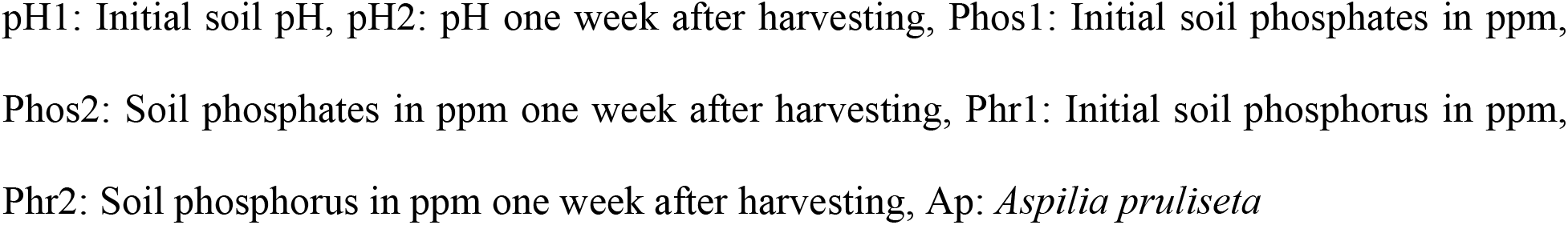
pH, phosphates and phosphorus content levels in soil before planting and one week after gadam sorghum crop harvest

| Factor | pH1 | pH2 | Phos1 | Phos2 | Phr1 | Phr2 |
| --- | --- | --- | --- | --- | --- | --- |
| **Silty Clay** | 6.3a | 6.2a | 96.2a | 96.8a | 25.7a | 24.2a |
| **Silt Loam** | 6.0b | 5.9b | 76.9a | 78.9a | 34.5a | 32.8a |
| **Sandy Loam** | 6.3a | 6.1a | 80.8a | 80.8a | 35.3a | 34.5a |
| **LSD** | 0.2 | 0.2 | 39.2 | 39.6 | 15.8 | 15.9 |
| **With Ap** | 6.0b | 5.9b | 103.6a | 107.7a | 31.2a | 30.0a |
| **Without Ap** | 6.3a | 6.2a | 65.6b | 63.3b | 31.8a | 31.0a |
| **LSD** | 0.2 | 0.2 | 32 | 32.3 | 12.9 | 13 |
| **0-20 cm** | 6.4a | 6.3a | 108.6a | 108.4a | 40.3a | 39a |
| **21-40 cm** | 6.1b | 5.9b | 65.3b | 67.3b | 28.5a | 27.8a |
| **41-60 cm** | 6.1b | 6.0b | 80.0ab | 80.8ba | 25.7a | 24.7a |
| **LSD** | 0.2 | 0.2 | 39.2 | 39.6 | 15.8 | 15.9 |
| **Mean** | 6.2 | 6.1 | 84.6 | 85.5 | 31.5 | 30.5 |

pH1: Initial soil pH, pH2: pH one week after harvesting, Phos1: Initial soil phosphates in ppm, Phos2: Soil phosphates in ppm one week after harvesting, Phr1: Initial soil phosphorus in ppm, Phr2: Soil phosphorus in ppm one week after harvesting, Ap: *Aspilium pruliseta*

## 4. Discussion

We found out that water retention capacity of the different soil types affected stand count with time. The smaller the particles were, the more the water holding capacity and with time, seeds that germinated dried up. [26] obtained similar results on pyrethrum seed emergence tested for different soil types while [27] observed that germination and emergence of *Helianthus annuus* L. was low in clay treatments compared to other soil treatments with bigger particle size. [28] varied rainfall, sowing time, soil type, and cultivar influence on plant population for wheat in Western Australia and obtained similar results.

Early growth of gadam sorghum plant was influenced by use of soils that had *Aspilia pruliseta* initially growing in it. [29] noted that mycorrhiza fungi negatively influenced seed germination but at the same time improved plant growth. This partly explains the behaviour of sorghum seedlings stand county stability compared to seedlings in soils that had not been growing *Aspilia pruliseta*.

Inoculation with mycorrhiza fungi had an effect on seed emergence, stand count and plant height in this experiment. Results obtained closely mirrored the observation made by [30] that rhizosphere aggregate stability of afforested semiarid plant species was significantly improved upon mycorrhiza fungi inoculation. Not only does mycorrhizal association with plants improve drought tolerance [31] but also seedling survival [32].

Gadam sorghum is a short stature crop [33] and this is confirmed in this experiment showing the plant as having an average height of 124 cm at physiological maturity. Having a small height is one of the drought escaping mechanism of plants in dry land areas as assimilate partitioning is done early during the plant’s life [34]

The study by [35] revealed that phosphorus was critical for cell division and elongation on the grass plants at early stages of growth. This phenomenon could have contributed towards greater plants height for seeds that were inoculated with mycorrhiza fungi in this current experiment and further corroborates research by [36] that Phosphorus inoculation was found to be positive and significant on mungbean plant height. In addition, [37] showed that germination and vigour in rice improved significantly when seeds were soaked in potassium and phosphorus salts.

Soil that had Ap (*Aspilia pruliseta*) previously growing, had mycorrhiza that acted as a source of inoculation and therefore continued with phosphates synthesis with gadam sorghum as the host plant. This finding agrees with [38] that mycorrhiza fungi inoculation enhances development of soil phosphates by increasing phosphates enzyme activity. The phosphorus in the soil was converted to phosphates through the action of mycorrhiza

## 5. Conclusion

Gadam sorghum plant emergence, stand count and aggregate plant stability is enhanced by inoculating seeds with Aspilia *pruliseta* rhizosphere soils. These attributes enhance overall crop yield and land productivity. Soils previously growing *Aspilia pruliseta* vegetation enhance phosphate amounts in the soil media and gadam sorghum is a good mycorrhiza host plant.

## Acknowledgement

Authors thank the National Research Fund in Kenya for the initial financial support and Agriculture Resource Management department team members in the University of Embu and Kenyatta University for reading and correcting this manuscript.

